# SRGAP2 limits experience-dependent structural synaptic plasticity in adult cortical circuits

**DOI:** 10.64898/2026.08.21.746251

**Authors:** Sergio Bernal-Garcia, Regina Jiang, Franck Polleux

## Abstract

In cortical circuits, synaptic plasticity involves either changes in the weight of pre-existing synapses, referred to as functional synaptic plasticity, or synapse formation and elimination, referred to as structural synaptic plasticity. Experience-dependent structural synaptic plasticity is prominent in juvenile cortical circuits during critical periods of development but drastically decreases in adult cortical circuits. The molecular mechanisms limiting experience-dependent structural synaptic plasticity in adult cortical circuits remain largely unknown. During development, the postsynaptic protein SRGAP2 limits the formation of both excitatory (E) and inhibitory (I) synapses in cortical pyramidal neurons (CPNs) and promotes their maturation. SRGAP2 expression is maintained throughout adulthood but its synaptic function in the adult cortex has not been explored. Using longitudinal 2-photon (2P) imaging of dendritic spine dynamics in layer 2/3 CPNs and found that this form of sensory deprivation induces a striking increase in structural synaptic plasticity favoring spine formation in adult constitutive SRGAP2^+/-^ mice, in contrast to wild-type adult mice, where whisker trimming does not induce significant structural synaptic plasticity. Using conditional, cell-type specific, deletion of SRGAP2, we demonstrate that this experience-dependent structural synaptic plasticity requires both of SRGAP2 in expression L2/3 CPNs and in microglia. We previously demonstrated that the human-specific paralogs SRGAP2B/C inhibit all known functions of SRGAP2, phenocopying SRGAP2 haploinsufficiency, our results suggest that SRGAP2B/C might endow increased levels of experience-dependent structural synaptic plasticity to human pyramidal neurons in adult cortical circuits.

## INTRODUCTION

In mammalian neocortical circuits, the modifiability of synapses underlies most forms of learning and memory either through (1) structural synaptic plasticity i.e. the elimination of existing synapses or the formation of new synapses, and/or (2) functional synaptic plasticity i.e. changes in synaptic weights of existing synapses. Whereas functional forms of synaptic plasticity are present in developing and adult cortical circuits (reviewed in ^1^), structural forms of synaptic plasticity are most pronounced during the juvenile postnatal period and declines steadily through adolescence to reach minimal levels in the adult cortex ^2,3^. For example, longitudinal *in vivo* two-photon (2P) structural imaging demonstrated that layer 2/3 (L2/3) pyramidal neurons show a marked reduction in baseline dendritic spine formation and elimination rates from adolescence into adulthood ^4–6^. This developmental reduction of dendritic spine dynamics is paralleled by a contraction of the structural plasticity response to changes in sensory experience: in the juvenile cortex, sensory deprivation drives extensive structural synaptic plasticity involving both increased spine formation and elimination ^7^ but in adult cortex, by contrast, sensory deprivation produces significantly more limited effects, primarily preventing spine elimination rather than driving new spine formation^8^. The molecular mechanisms that limit experience-dependent structural synaptic plasticity in adult cortical circuits remain largely unknown.

One of the most striking features characterizing human cortical neurons is their prolonged, neotenic, synaptic development spanning a decade in humans compared to months in other mammals including rodents and non-human primates. Synaptic neoteny in humans has been suggested to extend critical periods of cortical circuit plasticity underlying learning during postnatal development which might contribute to the emergence of advanced social, sensory and cognitive functions. Among the molecular candidates underlying neotenic features of synaptic development in human cortical neurons, the *Slit-Robo GTPase Activating Protein 2* (*SRGAP2;*^9^) gene family stands out. *SRGAP2* underwent a series of human-specific segmental gene duplications during the emergence of the *Homo* lineage, generating two partial gene duplications (*SRGAP2B*, *SRGAP2C*) expressing truncated proteins that are highly unstable structurally ^10^ and are constitutively targeted to the proteosome for degradation ^11^ but remain able to heterodimerize with the ancestral full-length SRGAP2A protein and decrease by ∼50% of its abundance in cortical neurons ^12,13^. Humanization of SRGAP2C expression in mouse cortical neurons therefore phenocopies a partial loss of function of SRGAP2, increasing the density of both excitatory (E) and inhibitory (I) synapses on L2/3 pyramidal neurons while preserving E/I balance, and extending the timing of synapse maturation ^12,14^. These changes are collectively described as synaptic neoteny: a retention of juvenile-like features of synaptic maturation into adulthood. SRGAP2A normally drives the developmental closure of these programs, such that its inhibition by SRGAP2C, or its partial loss in SRGAP2^+/-^ mice, prolongs the timing of their maturation ^11–15^. At the circuit and behavioral level, SRGAP2C expression in mouse cortical pyramidal neurons increases local and long-range cortico-cortical connectivity, increases the reliability of sensory coding, and improves learning of a complex sensory discrimination task compared to wild-type mice ^15^.

Interestingly, *SRGAP2* and its human-specific paralogs *SRGAP2B/C* are not only expressed in developing and adult cortical neurons but are also expressed at high levels in microglial cells (see **Figure 1** and ^16^). Conditional deletion of *Srgap2* specifically in microglia induces developmental neoteny of their structural and functional microglia maturation which contributes, in a non-cell autonomous manner, to synaptic neoteny of cortical pyramidal neurons and contributes to the net increase in spine density in adult layer 2/3 PNs ^16^.

**Figure 1.**
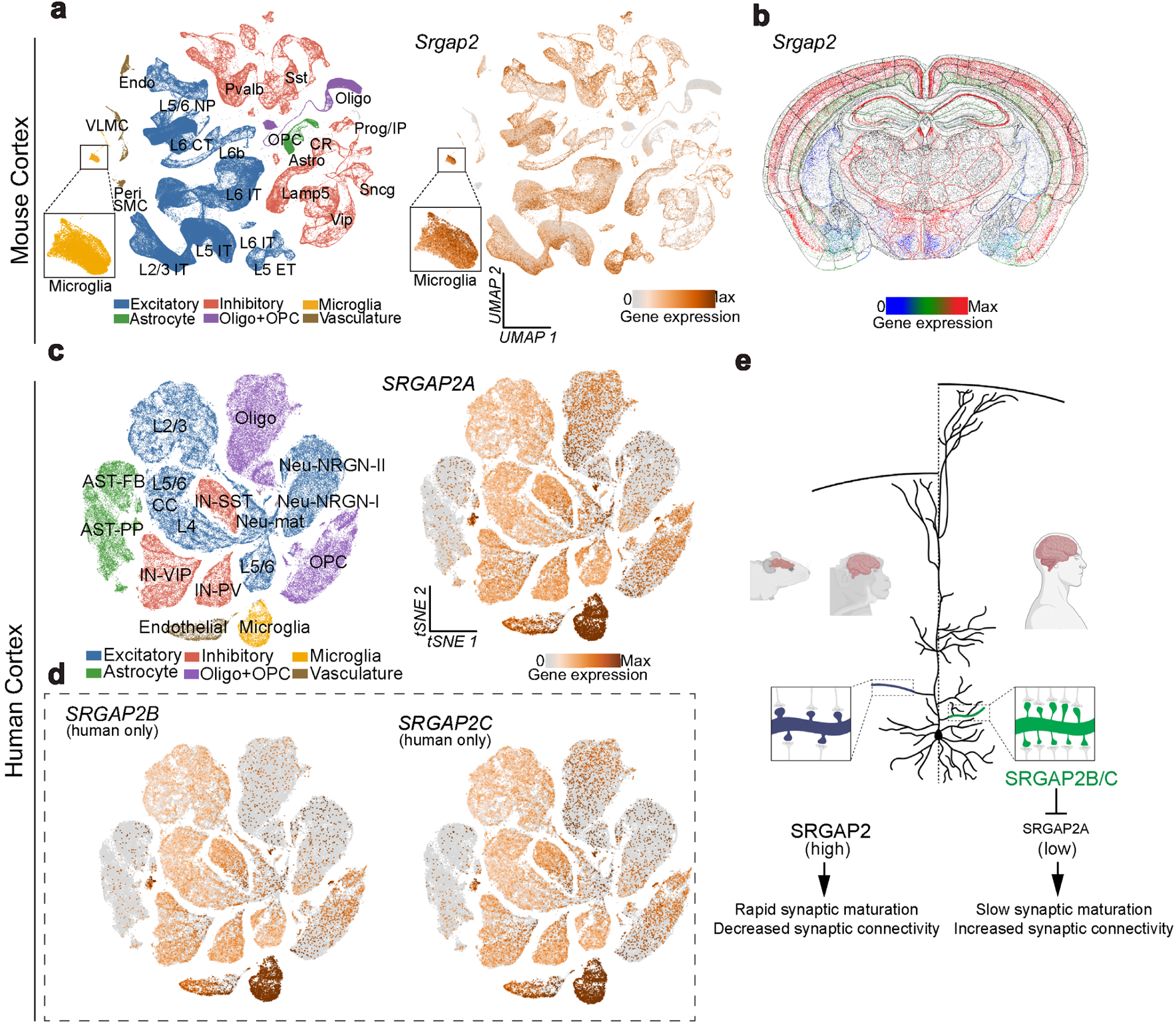
Expression of *SRGAP2A* and its human-specific paralogs *SRGAP2B/C* in the adult cortex. **a** | Single-cell transcriptomes of the adult mouse motor cortex (UCSC Cell Browser, *mouse-mop-atlas*). Left, UMAP colored by cell class: Excitatory (blue; subclasses L2/3 IT, L5 IT, L5 ET, L5/6 NP, L6 CT, L6 IT, L6 IT Car3, L6b), Inhibitory (red; Pvalb, Sst, Vip, Lamp5, Sncg), Microglia (orange), Astrocyte (green; Astro), Oligo+OPC (purple; Oligo, OPC), Vasculature (brown; Endo, VLMC, SMC, Peri), and Other (gray; CR, Prog/IP). Right, the same UMAP colored by *Srgap2* expression, shown as discrete transcript-count bins (0, 1, 2, …, 17+) on a sequential color scale from light gray (zero) through orange to dark brown (highest bin). **b** | Imputed spatial expression of *Srgap2* on a representative coronal section of the adult mouse brain (MERFISH-based imputation, Allen Brain Cell Atlas). Color encodes relative gene expression from low (blue) through intermediate (green) to high (red). **C** | Single-cell transcriptomes of the developing and adult human cortex (UCSC Cell Browser, *autism* dataset). Left, tSNE colored by cell class (palette as in **a**): Excitatory (blue; L2/3, L4, L5/6, L5/6-CC, Neu-NRGN-I, Neu-NRGN-II, Neu-mat), Inhibitory (red; IN-PV, IN-SST, IN-SV2C, IN-VIP), Astrocyte (green; AST-FB, AST-PP), Microglia (orange), Oligo+OPC (purple; Oligodendrocytes, OPC), and Vasculature (brown; Endothelial). Right, the same tSNE colored by *SRGAP2A* expression on the same gray-to-brown color scale as in **a**, discretized into 11 bins (zero plus ten quantile bins of nonzero normalized expression). **d** | The human tSNE shown in **c**, colored by expression of the human-specific paralogs *SRGAP2B* (left) and *SRGAP2C* (right), each binned and colored as in **c**. Dashed box indicates that these paralogs are only present in the human genome. **e** | Schematic of *SRGAP2* paralog function. In mouse (*Srgap2*) and non-human primates (*SRGAP2*) (left), only the ancestral *SRGAP2* gene is expressed, driving rapid synaptic maturation and decreased synaptic connectivity. In humans (right), the duplicated paralogs SRGAP2B and SRGAP2C heterodimerize with and inhibit SRGAP2A, slowing synaptic maturation and increasing synaptic connectivity. Insets compare spine density on a representative cortical pyramidal neuron under high ancestral SRGAP2 activity (blue, left; mouse/non-human primates) versus SRGAP2A inhibited by SRGAP2B/C (green, right; human).

Since (1) *SRGAP2* and its human-specific paralogs *SRGAP2B/C* are expressed in adult neurons and microglia and (2) that during development, SRGAP2 functions by limiting dendritic spine formation (see summary in **Fig. 1e**; ^11–15^), we hypothesized that adult expression of SRGAP2 could play a role in limiting the expression of structural synaptic plasticity. We tested this using *in vivo* 2-photon structural and longitudinal imaging of spine dynamics in both constitutive SRGAP2^+/-^ knockout mice as well as conditional, cell-type specific, conditional deletion of SRGAP2 in L2/3 PNs or microglia. Our results demonstrate that SRGAP2 constitutive haploinsufficiency leads to increased rates of spine formation (but not elimination) in adult L2/3 PNs compared to wild-type mouse control. Strikingly, while whisker trimming does not increase spine formation in adult wild-type cortex as previously shown ^2,3,8^, we observed significant increase in spine turnover, dominated by increased spine formation, in adult SRGAP2^+/-^ mice. Interestingly, our analysis of spine dynamics following sensory deprivation using cell-type specific, conditional, deletion of SRGAP2 in L2/3 PNs or microglia reveal that SRGAP2 expression in both cell types contributes to the limitation of experience-dependent structural synaptic plasticity in adult cortex. Since the expression of human-specific paralogs *SRGAP2B/C* is necessary and sufficient to decrease SRGAP2 by ∼50% at the protein level ^12,13^, our new results raise the intriguing possibility that human CPNs might display heightened levels of experience-dependent structural synaptic plasticity in adult cortical circuits, a trait that might have contributed to the emergence of increased learning and cognitive capacities during human evolution.

## RESULTS

### *Srgap2* and its human-specific paralogs are expressed in cortical pyramidal neurons and in microglial cells in the adult cortex

We have previously demonstrated that SRGAP2 and its human-specific paralogs are expressed throughout development in cortical pyramidal neurons ^12,13,17^ and in microglia ^16^. However, *SRGAP2* and the human-specific *SRGAP2B/C* paralogs are also expressed in the adult cortex (**Fig. 1c-d**). We leveraged single cell/nucleus RNA sequencing (snRNAseq) data from publicly accessible resources to determine the cell types expressing *SRGAP2* in the adult mouse motor cortex ^18^. *SRGAP2* is expressed by all classes of cortical neurons including L2/3 PNs as well as by microglial cells (**Fig. 1a**). This was confirmed by examination of spatial transcriptomics data ^19^ from the adult mouse cortex where *SRGAP2* mRNA is detected at high level throughout the adult cortex and at highest levels in layer 2/3 (**Fig. 1b**). Using publicly accessible snRNAseq data from human cortex^20^, we confirmed that both ancestral *SRGAP2A* and human-specific paralogs *SRGAP2B/C* mRNA are detected in all classes of cortical neurons including L2/3 PNs as well as in high expression in microglia (**Fig. 1c-d**).

### Longitudinal *in vivo* 2P imaging of L2/3 pyramidal-neuron spines in adult barrel cortex

To determine whether SRGAP2 plays a role in the expression of structural synaptic plasticity in L2/3PNs of the adult cortex, we established a longitudinal *in vivo* two-photon (2P) imaging paradigm in order to track individual dendritic spine dynamics across eight consecutive daily sessions in sparsely labeled L2/3 PNs of the adult mouse barrel cortex (**Fig. 2a**).

**Figure 2.**
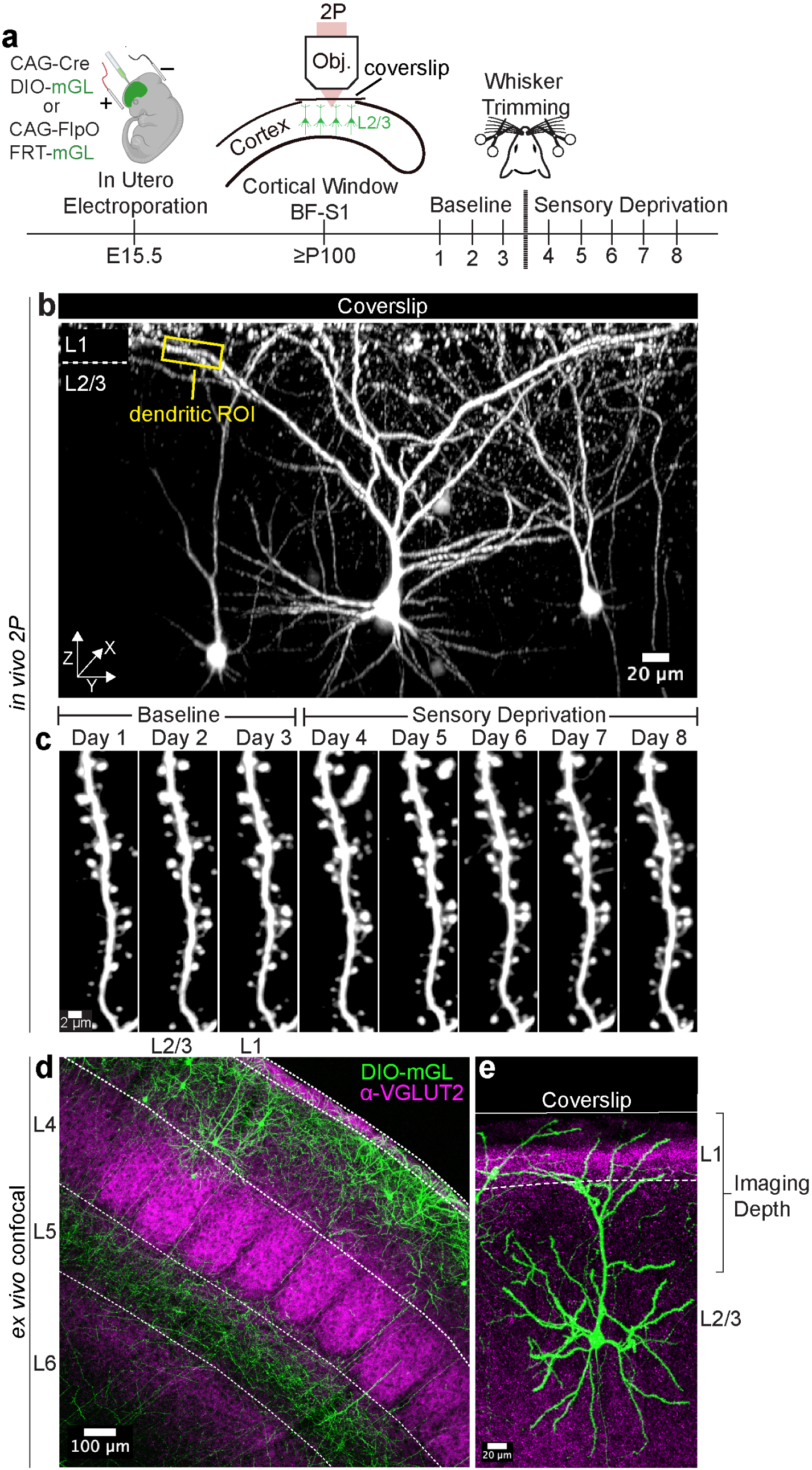
Longitudinal *in vivo* two-photon imaging of sparsely labeled layer 2/3 pyramidal neurons in the adult barrel cortex. (**a**) Experimental design. Layer 2/3 pyramidal neurons in barrel field somatosensory cortex (BF-S1) were sparsely labeled via *in utero* electroporation (IUE) at embryonic day 15.5 (E15.5) using a dual-recombinase strategy (CAG-Cre/DIO-mGreenLantern and CAG-FlpO/FRT-mGreenLantern). A chronic cranial window was implanted over BF-S1 at or after postnatal day 100 (P100). Dendritic spines were imaged daily with two-photon (2P) microscopy across 8 consecutive sessions. Baseline imaging was performed from Days 1-3; sensory deprivation was induced by bilateral whisker trimming beginning after session 3; sensory deprivation imaging was done from Days 4-8. (**b**) Representative *in vivo* 2P maximum intensity projection of a sparsely labeled L2/3 pyramidal neuron, showing the full apical dendritic arbor, soma and basal dendrites extending from L2/3 into L1 beneath the coverslip. A dendritic region of interest (ROI, yellow) selected for longitudinal spine tracking is indicated. Scale bar, 20 μm. **c** | Daily time-lapse images of a single dendritic segment across all 8 sessions (Day 1 to Day 8), demonstrating the resolution and stability of chronic imaging for tracking individual spines. Scale bar, 2 μm. (**d**) *Ex vivo* confocal image of a coronal section through the barrel cortex showing the laminar distribution of electroporated neurons. Sparse L2/3 pyramidal neurons express mGreenLantern (green). vGluT2 immunostaining (magenta, far-red secondary antibody) demarcates the barrels in L4 beneath the cortical window. Cortical layers (L1 to L6) are indicated. Scale bar, 100 μm. (**e**) Higher-magnification *ex vivo* confocal image of individual L2/3 pyramidal neurons (green), showing complete apical dendritic arbors within the imaging depth accessible through the cranial window. Scale bar, 20 μm.

L2/3 PNs located in the barrel field of the primary somatosensory cortex (BF-S1) were sparsely labeled with bright fluorescent protein (mGreenLantern; mGL) by *in utero* electroporation (IUE) at embryonic day (E) 15.5, using a dual-recombinase strategy that independently controlled labeling density and fluorophore intensity (**Fig. 2a**). Low amount of plasmids expressing Cre or FlpO recombinase were co-electroporated with Cre- or FlpO-dependent mGL expressing plasmids (DIO-mGreenLantern for Cre or FRT-mGreenLantern for FlpO), yielding sparse expression of mGL in L2/3 neurons. At postnatal day 100 (P100), a chronic cranial window was implanted stereotactically over the BF-S1 cortex, and daily 2P imaging was performed across eight consecutive sessions. This approach enabled longitudinal imaging of the same individual, optically isolated, L2/3PNs, their dendrites and dendritic spines for eight continuous days spanning baseline (days 1-3) and following sensory deprivation by whisker trimming (days 4-8) (**Fig. 2a-c**).

Our structural *in vivo* imaging revealed the entire neuronal architecture of sparsely labeled neurons from deep basal dendrites in L2/3 through the apical trunk and distal tuft extending into L1 beneath the cranial window (**Fig. 2b and e**). Bright and optically isolated distal tuft dendritic segments directly under the cortical window were selected at Day 1 and relocated in each subsequent session, allowing individual spines to be tracked across all eight sessions with preserved signal-to-noise and individual spine resolution (**Fig. 2c**). Post hoc confocal imaging of fixed coronal sections confirmed the spatial and laminar specificity of the IUE cohort: mGL-expressing neurons were confined to L2/3 of BF-S1 and aligned directly above the VGluT2-demarcated barrels in L4 that lay beneath the cranial window (**Fig. 2d**), and their apical dendritic arbors fell entirely within the imaging depth accessible through the window (**Fig. 2e**). Across all genotypes analyzed in this study (wild-type (WT) controls, *Srgap2*^+/-^, L2/3-*Srgap2*^F/+^, and MG-*Srgap2*^F/F^; see **Fig. 3a**), this paradigm yielded a longitudinal dataset of 22,420 individual dendritic spines tracked at single-spine resolution across eight consecutive daily sessions (264 branches, 19 mice, 56,728 spine-pair observations spanning pre-deprivation baseline and sensory deprivation; see Supplemental Table 1).

**Figure 3.**
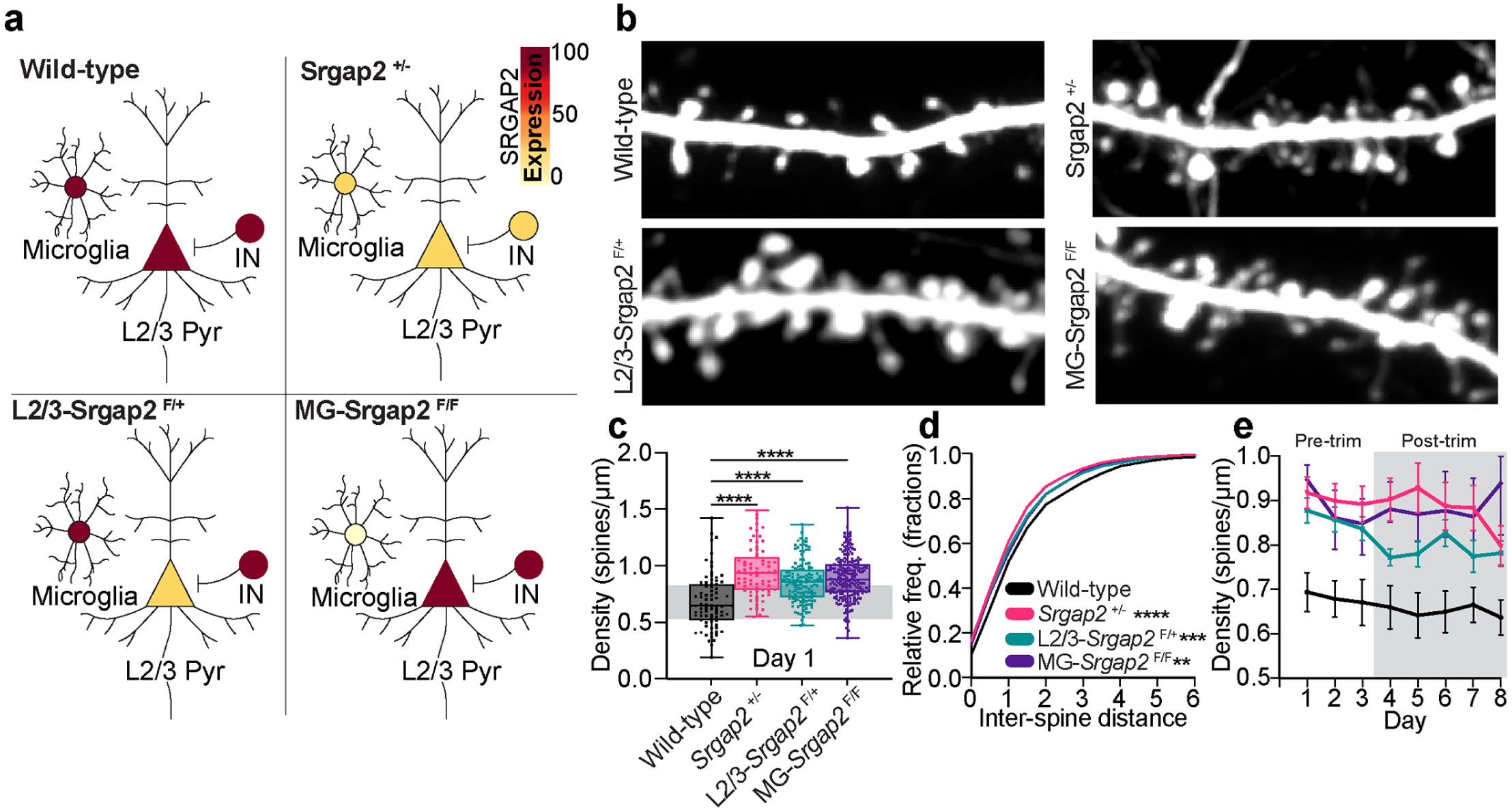
Cell-autonomous and non-cell-autonomous *Srgap2* reduction stably increases spine density in adult L2/3 cortical pyramidal neurons. (**a**) Schematic of genotypes and SRGAP2 protein expression levels across the three major cell types manipulated in this study. Wild-type: 100% SRGAP2 expression in L2/3 pyramidal neurons (L2/3 Pyr), microglia, and interneurons (IN). *Srgap2*^+/-^: ∼50% expression in all cell types (global heterozygous). L2/3-*Srgap2*^F/+^: ∼50% expression cell-autonomous to targeted L2/3 pyramidal neurons. MG-*Srgap2*^F/F^: near-zero expression of SRGAP2 in microglia only. (**b**) Representative Day 1 maximum intensity projections of dendritic segments for each genotype. Scale bar, 2 μm. (**c)** Baseline spine density (Day 1) across genotypes. Each point represents an individual dendritic segment. One-way ANOVA against Wild-type. ****P < 0.0001. (**d**) Cumulative distribution of inter-spine distances at Day 1 for each genotype. Wild-type (black), *Srgap2*^+/-^ (red), L2/3-*Srgap2*^F/+^ (teal), MG-*Srgap2*^F/F^ (purple). Kolmogorov-Smirnov test against Wild-type. (**e**) Longitudinal spine density across Days 1-8 for the same dendritic segments tracked over time. Pre-trim (Days 1-3) and Post-trim (Days 4-8) epochs are indicated. Colors as in (d). Error bars, s.e.m. n = 19 mice, 264 branches, 22,420 spines. **P < 0.01, ***P < 0.001, ****P < 0.0001.

To track individual dendritic spines across eight imaging sessions, four genotypes, and two labeling strategies, we developed a custom analysis pipeline coupling our recently development RESPAN framework for spine segmentation ^21^ with cross-session volumetric registration (via itk-elastix), 3D label refinement, and longitudinal Hungarian-algorithm matching (see Methods; **Suppl. Fig. 1**). Registration increased the median normalized cross-correlation (NCC, a volumetric similarity score ranging from 0 to 1) between registered moving and Day 1 reference volumes from approximately 0.31 to approximately 0.80 across the full dataset (**Suppl. Fig. 1c,d**). The median residual centroid displacement between consecutive registered sessions for matched spines was 0.499 µm, with 92.6% of matched-spine displacements falling below 1.0 µm (approximately the diameter of a single spine head), supporting confident cross-session spine identity assignment (**Suppl. Fig. 1e**).

### Cell-autonomous and non-cell-autonomous *Srgap2* reduction stably increases spine density in adult L2/3 pyramidal neurons

We previously shown that constitutive heterozygous removal of *Srgap2* (*Srgap2*^+/-^) leads to a developmental increase in spine density compared to WT control littermates that remains elevated into adulthood (>P70) ^11–14^. To test whether removal of *Srgap2* modulates experience-dependent dendritic spine dynamics in adult cortical circuits, we analyzed three complementary cohorts in which *Srgap2* expression was reduced across distinct cell-types (**Fig. 3a**): (1) a constitutive global heterozygous knockout (*Srgap2*^+/-^), which reduces SRGAP2 protein expression by ∼50% in every cell type expressing SRGAP2 ^12^, (2) a conditional (Floxed allele; *Srgap2*^F/+^) heterozygous line we previously characterized ^22^ in which *Srgap2* deletion is restricted cell-autonomously to the Cre-expressing, sparsely electroporated L2/3 pyramidal neurons, and (3) a microglia-specific conditional homozygous knockout (MG-*Srgap2*^F/F^) in which *Srgap2* is inducibly deleted in microglia via a tamoxifen-inducible Cre^ERT2^ under the microglia-specific TMEM119 promoter (induced by daily tamoxifen administration from postnatal days 3-5). In the latter case, we sparsely labeled L2/3 PNs using CAG-FlpO and FRT-mGreenLantern, therefore maintaining wild-type *Srgap2* expression in L2/3 PNs (**Fig. 3a**). Together, these three genotypes allowed us to dissect the contributions of *Srgap2* dosage when haploinsufficiency is achieved in all cells normally expressing SRGAP2 (*Srgap2*^+/-^), when haploinsufficiency is achieved in L2/3 PNs cell-autonomously (L2/3-*Srgap2*^F/+),^ or when *Srgap2* is bi-allelically deleted in microglia (MG-*Srgap2*^F/F^), isolating the non-cell-autonomous contribution of microglial SRGAP2 to adult L2/3 PN spine density (**Fig. 3**) and spine dynamics (**Fig. 4**).

**Figure 4.**
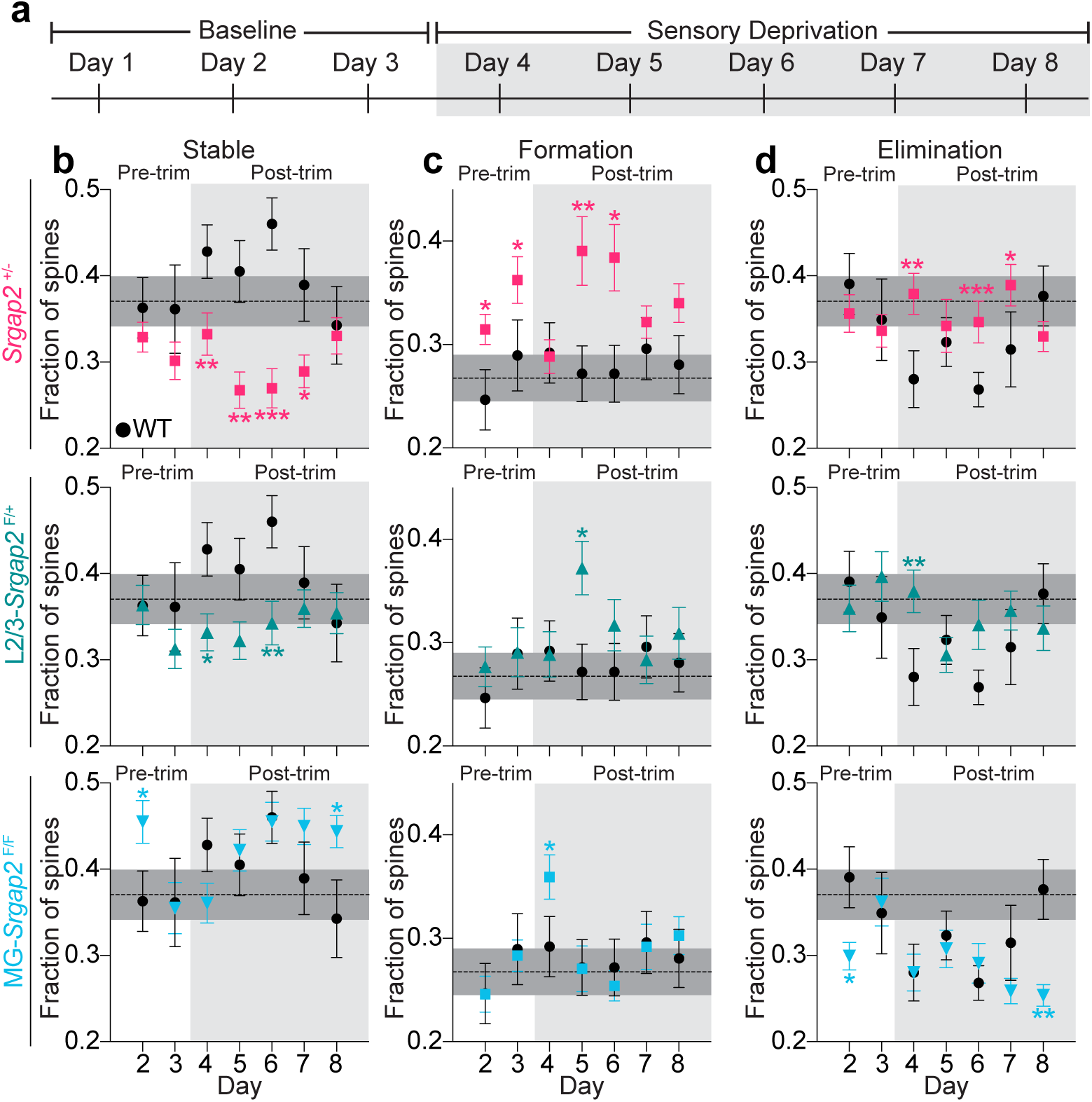
Cell-autonomous *Srgap2* reduction in L2/3 pyramidal neurons permits experience-dependent spine dynamics in the adult cortex. (**a)** Experimental timeline. Baseline imaging on Days 1-3 (white) and sensory deprivation imaging on Days 4-8 (gray shading), with bilateral whisker trimming initiated after Day 3. (**b-d**) Fraction of spines classified as Stable (b), Formation (newly formed, c), and Elimination (eliminated, d) across consecutive day pairs (x-axis: the later day of each pair; e.g., Day 2 = pair Days 1-2). Each column plots three mutant genotypes in separate rows against WT (black circles, all rows): *Srgap2*^+/-^ (pink squares, top), L2/3-*Srgap2*^F/+^ (teal triangles, middle), and MG-*Srgap2*^F/F^ (cyan downward triangles, bottom). Pre-trim (Days 1-3, white) and post-trim (Days 4-8, gray shading) days are indicated. The horizontal dashed line and surrounding gray band indicate the WT baseline mean (Day pairs 1-2 and 2-3) ± s.e.m. for the corresponding category. Two-sided Mann-Whitney U test, each genotype × day compared against WT on the same day. Error bars, s.e.m. per dendritic branch. n = 19 mice, 264 branches, 22,420 spines. *P < 0.05, **P < 0.01, ***P < 0.001.

We validated that our approach to induce SRGAP2 conditional deletion specifically in microglia using the TMEM119-Cre^ERT2^;SRGAP2^F/F^ approach following tamoxifen induction at P3-5 leads to microglia hyper-ramification that is sustained in the adult cortex (**Fig. S2**) as recently described during development ^16^.

At the pre-sensory deprivation baseline (Day 1), each of the three *Srgap2* loss-of-function (LOF) genotypes displayed significantly higher spine density along apical dendritic segments compared to Wild-type (**Fig. 3b**). Quantitatively, Day 1 spine density was significantly elevated in *Srgap2*^+/-^ (0.951 ± 0.026 spines/µm), L2/3-*Srgap2*^F/+^ (0.857 ± 0.016 spines/µm), and MG-*Srgap2*^F/F^ (0.892 ± 0.013 spines/µm) relative to wild-type control (0.680 ± 0.027 spines/µm) (Kruskal-Wallis test with Dunn’s post-hoc comparisons against WT controls: *Srgap2*^+/-^, Z = 7.08, p < 0.0001; L2/3-*Srgap2*^F/+^, Z = 5.37, p < 0.0001; MG-*Srgap2*^F/F^, Z = 7.26, p < 0.0001; **Fig. 3c**). Consistent with elevated spine density, the cumulative distribution of inter-spine distances on Day 1 dendritic segments was significantly left-shifted in all three mutant genotypes relative to WT (two-sample Kolmogorov-Smirnov test against Wild-type: *Srgap2*^+/-^, D = 0.103, p < 0.0001; L2/3-*Srgap2*^F/+^, D = 0.082, p = 0.0005; MG-*Srgap2*^F/F^, D = 0.071, p = 0.0016; **Fig. 3d**), indicating that the increase in spine density reflected a generalized reduction in the spacing between adjacent spines along dendritic segments. Notably, we observed a genotype-dependent decrease in the cumulative distribution of inter-spine distances, which is a more sensitive way to quantify changes in spine density compared to computing spine density per dendritic segments: *Srgap2*^+/-^ showed the strongest left shift in the distribution (shortest inter-spine distance), while L2/3-*Srgap2*^F/+^ and MG-*Srgap2*^F/F^ inter-spine distances displaying strikingly intermediate distributions to those of *Srgap2*^+/-^ and WT. Therefore, our quantification reveals that the increased spine density characterizing L2/3PNs in the adult constitutive *Srgap2*+/- heterozygous mice is the combined result of cell autonomous SRGAP2 expression in L2/3PNs as well as a cell non-autonomous contribution through SRGAP2 expression in microglia ^16^.

The elevated spine density in *Srgap2*-LOF genotypes was maintained across all eight imaging sessions: density in each mutant genotype remained well above the Wild-type trajectory throughout both the pre-deprivation baseline (Days 1-3) and the sensory deprivation period (Days 4-8), with no detectable decrease toward Wild-type levels and no trending increase across the deprivation period (**Fig. 3e**).This lack of a change in spine density following sensory deprivation held across all four genotypes, including the MG-*Srgap2*^F/F^ cohort, in which conditional deletion of *Srgap2* in microglia did not detectably modulate spine density during deprivation despite the canonical role of microglia in activity-dependent synaptic remodeling ^23–25^. Spine density therefore did not exhibit a clear sensory deprivation-dependent change in any of SRGAP2 genotypes examined. However, spine density is an inherently coarse metric of structural synaptic plasticity since the same spine density might be observed regardless of magnitude of dendritic spine dynamics. Indeed, prior longitudinal *in vivo* imaging studies of adult cortex have repeatedly shown that experience-dependent changes in spine turnover can occur without detectable changes in spine density ^3,8^.

### *Srgap2* limits experience-dependent spine formation in the adult cortex through its expression in both L2/3 pyramidal neurons and microglia

To test whether *Srgap2* plays a role in the expression of experience-dependent structural synaptic plasticity in adult L2/3 PNs, we quantified the fraction of stable, newly formed, and eliminated spines along the same dendritic segments before and after bilateral whisker trimming. We plotted each *Srgap2*-LOF genotypes against WT controls on consecutive day pairs across the pre-sensory deprivation baseline (Day pairs 1-2 and 2-3) and the post whisker trimming (Day pairs 3-4 through 7-8) (**Fig. 4a**).

In WT control mice, sensory deprivation increased the fraction of stable spines above the pre-trim baseline, the fraction of eliminated spines decreased, and the fraction of newly formed spines remained at its pre-trimming level (**Fig. 4b-d**, black circles, all rows). By contrast, in constitutive *Srgap2*^+/-^, we observe a significant reduction in the fraction of stable spines relative to WT, particularly during the post-deprivation window. In *Srgap2*^+/-^, this is due to a significant increase in the rate of spine formation compared to WT following sensory deprivation at multiple post-trim days, with the strongest differences on Day pairs 4-5 (0.39 vs 0.27, p = 0.002) and 5-6 (0.38 vs 0.27, p = 0.033) (**Fig. 4c**, first row).

In dendrites of L2/3-*Srgap2*^F/+^ neurons, in which *Srgap2* heterozygosity is restricted cell-autonomously to the imaged L2/3 pyramidal neurons, exhibited a significant reduction of the fraction of stable spines compared to WT controls but a significant increase in the rate of spine formation at Day pair 4-5 (0.37 vs 0.27, p = 0.041) without a detectable elevation at pre-trim baseline (**Fig. 4c**, middle row). The rate of spine elimination in dendrites of L2/3-*Srgap2*^F/+^ remain stable following whisker trimming which is significantly different from WT controls when spine elimination decreases following sensory deprivation (**Fig. 4d**, top and middle rows).

Following microglial-restricted deletion of SRGAP2 (MG-*Srgap2*^F/F^), L2/3 PNs display slight but significant changes in the fraction of stable spines but a significant increase in the rate of spine formation one day after whisker trimming compared to WT (day 4) and an even more pronounced decrease in the rate of spine elimination following sensory deprivation (day 8) compared to WT (**Fig. 4b, d**, bottom rows).

We were next interested in understanding whether SRGAP2 altered the spine size of the stable or the transient population of spines. For every consecutive day pair, we measured the area of each spine and compared the stable population with the transient population of newly formed and eliminated spines (**Suppl. Fig. 3**). In all four genotypes, stable spines were approximately twofold larger than transient spines, consistent with the established relationship between spine size and stability and confirming that our segmentation and tracking pipeline faithfully recovered this relationship across sessions ^5,26^ (**Suppl. Fig. 3a-h**). Relative to wild-type, the stable spine population was significantly enlarged in *Srgap2*^+/-^ (median spine area 0.749 vs 0.697 µm^2^; Kolmogorov-Smirnov D = 0.069, p = 0.003) and in MG-*Srgap2*^F/F^ (0.801 vs 0.697 µm^2^; D = 0.080, p = 0.0003), but was indistinguishable from wild-type in L2/3-*Srgap2*^F/+^ (0.697 vs 0.697 µm^2^; n.s.) (**Suppl. Fig. 3f-h**). The size of transient spines did not differ from wild-type in any genotype. Enlargement of the stable spine population therefore appeared selectively when SRGAP2 was reduced in microglia, in *Srgap2*^+/-^ and MG-*Srgap2*^F/F^, and was absent when SRGAP2 was reduced cell-autonomously in L2/3 PNs alone, implicating microglial SRGAP2 as a non-cell-autonomous regulator of stable spine size.

Taken together, these results demonstrate that (1) SRGAP2 limits the expression of experience-dependent structural synaptic plasticity, specifically limiting the rate of spine formation following sensory deprivation in adult layer 2/3 pyramidal neurons and (2) that this function of SRGAP2 is mediated through combined expression in both L2/3 PNs and microglia.

## DISCUSSION

The molecular mechanisms limiting the structural synaptic plasticity in adult cortical circuits remains largely unknown. Here, we tracked dendritic spine dynamics by longitudinal *in vivo* two-photon imaging of layer 2/3 pyramidal neurons in the adult mouse barrel cortex, across genotypes in which SRGAP2 was reduced in different cell types. Each of the three *Srgap2* loss-of-function genotypes carried an elevated spine density that remained stable across imaging and unchanged by sensory deprivation (**Fig. 3**); the response to sensory deprivation appeared not in density but in the dynamics of individual spines (**Fig. 4**). In wild-type adult mouse cortex, whisker trimming shifted L2/3 pyramidal neurons toward structural stability by reducing dendritic spine elimination, leaving formation largely unchanged. By contrast, reducing SRGAP2, whether constitutively in all cell types expressing SRGAP2 (*Srgap2*+/-), cell-autonomously in L2/3 pyramidal neurons (L2/3-*Srgap2*^F/+^), or cell-autonomously in microglia (MG-*Srgap2*^F/F^), unmasked an experience-dependent increase in spine formation absent in wild-type adults and, in *Srgap2*^+/-^ mice, already evident at baseline. Because SRGAP2 and its human-specific paralogs are expressed in both pyramidal neurons and microglia in the adult cortex (**Fig. 1**), these results demonstrate that SRGAP2 actively suppresses experience-dependent structural synaptic plasticity in the adult cortex, acting both cell-autonomously within L2/3 pyramidal neurons and non-cell-autonomously in microglia.

Spine formation is the structural substrate of synaptic plasticity, the process by which a neuron samples presynaptic partners from the large pool of surrounding axons as new presynaptic partners ^3,27–29^. In the adult cortex, experience-dependent structural synaptic plasticity is limited, and our results identify SRGAP2 as one of the molecular constraints enforcing that suppression. Reducing SRGAP2 partially reopens the capacity of L2/3 pyramidal neurons to add spines in response to reduction of sensory experience, a quantitative shift toward the elevated turnover that characterizes the juvenile cortex^5,6,8^ rather than a literal recapitulation of the juvenile state, in which sensory deprivation acts predominantly by slowing elimination ^8^. Longitudinal imaging has established that the baseline dynamics and experience-dependent plasticity of dendritic spines differ markedly between pyramidal neurons of different cortical layers ^6,30^. Because SRGAP2 is also expressed by layer 5 PNs (**Fig. 1a**), an important next step will be to determine whether its loss unmasks a comparable increase in experience-dependent spine formation in L5 PNs, or whether the constraint that SRGAP2 imposes on structural plasticity operates differently across PNs in different cortical layers.

SRGAP2 haploinsufficiency (*Srgap2*^+/-^) reveals that this expanded capacity is not confined to the deprivation period. The rate of spine formation is elevated above wild-type levels at baseline, prior to sensory deprivation, and was further increased toward spine formation following whisker trimming (**Fig. 4**). Since SRGAP2 is a postsynaptic protein ^12^, these changes are confined to spines and leave dendritic and axonal branch architecture intact, as is characteristic of adult cortical structural plasticity ^3^. A comparable reduction of SRGAP2, modeled by humanized expression of SRGAP2C in all cortical pyramidal neurons, increases cortico-cortical connections received by layer 2/3 pyramidal neurons, specifically the local feedforward inputs from layer 4 and long-range feedback inputs ^15^. These humanized SRGAP2C mouse model also improves the ability of these mice to learn a complex, whisker-based, sensory discrimination task ^15^. We attributed these changes in behavioral performance to the increased cortico-cortical connectivity, but future experiments will need to explore if the heightened structural synaptic plasticity we report in this study also contribute to improve learning in this mouse model, since structural synaptic plasticity has been shown to underlie some forms of learning.

The molecular mechanisms underlying the ability of SRGAP2 to limit experience-dependent structural synaptic plasticity remains unexplored. SRGAP2 contains a Rac1-specific GAP domain promoting GTP hydrolysis of Rac1 and therefore inactivating Rac1 ^31^. Rac1 is a small GTPase playing crucial roles in spine morphogenesis, synaptic maturation and structural and functional forms of synaptic plasticity (reviewed in^32^). Rac1 expression in neurons increases throughout synaptogenesis, and its early ectopic expression drives spine formation and AMPAR recruitment thereto ^33^. Increasing Rac1 activation, either through the activation of the Rac1-GEFs Kalirin-7 ^34,35^, β-PIX ^36^ or Tiam1 ^37,38^, or the knockout or knockdown of the Rac1-GAPs Bcr/Abr^39^, RICH2/ARHGAP44 ^40^, or p250GAP ^41^ drives synaptogenesis and structural synaptic plasticity during postnatal development. On the other hand, suppressing Rac1 activation using dominant negative Rac1 ^42^, Rac1-GAP overexpression ^39,43^ decreases spine density and synaptic plasticity.

Furthermore, we recently uncovered that, during synaptic maturation, SRGAP2, which promotes synaptic maturation, functions by cross-inhibiting the function of another postsynaptic protein SYNGAP1, a Ras-specific GAP protein. Beyond the well-documented function of SYNGAP1 as a negative regulator of excitatory synaptic maturation ^44,45^, SYNGAP1 has been best characterized for its role in functional and structural synaptic plasticity (reviewed in ^46^). Therefore, the antagonism between SRGAP2A and SYNGAP1 offers the most direct candidate molecular mechanism. SRGAP2A and SYNGAP1 are reciprocally inhibiting their own accumulation postsynaptically and are expressed in a mutually exclusive manner at the level of individual spines, where they exert opposite effects on the timing of synapse formation and maturation ^13^. SRGAP2 haploinsufficiency essentially represent a SYNGAP1 gain-of-function. Haploinsufficiency of Syngap1 accelerates the structural maturation of cortical pyramidal neurons, reduces dendritic spine turnover, and abolishes the increase in dendritic spines dynamics induced by sensory deprivation cortex ^44^}. In xenotransplanted human cortical neurons, SYNGAP1 loss of function likewise reduced dendritic spine turnover, a premature stabilization that disrupts their normally protracted, neotenic synaptic maturation ^13,45^. This reduced turnover is the direct mirror image of the increased spine turnover and formation we observed under SRGAP2 haploinsufficiency, consistent with a gain of SYNGAP1 function. Future experiments will need to test if the phenotypes observed in SRGAP2+/- mice showing increased experience-dependent spine turnover are suppressed by removing one copy of SYNGAP1 using SRGAP2^+/-^;SYNGAP1^+/-^ mice.

Newly formed spines acquire the postsynaptic scaffold PSD95 only as they stabilize, and those that fail to recruit it remain small and transient and are eliminated within days, accounting for the majority of spine turnover in the adult neocortex ^47,48^. It is this recruitment step, and not the formation of the spine itself, that is experience-dependent, with activity driving PSD95 accumulation at nascent spines through the activity-regulated protein CPG15/Neuritin while new spines continue to form largely independently of sensory experience^49^.

Cortical spines also carry an inhibitory dimension that SRGAP2 is well placed to control. A subset of dendritic spines are dually innervated, receiving an inhibitory synapse alongside their excitatory one, and on these spines the inhibitory synapse is far more dynamic than the spine itself, with gephyrin puncta repeatedly assembled and removed while the underlying spine remains structurally stable ^50,51^. SRGAP2A binds gephyrin through its C-terminal SH3 domain and organizes the inhibitory postsynaptic scaffolding of GABA-A receptors^14^. Its reduction may therefore reshape the assembly and removal of gephyrin on dually innervated spines, an inhibitory dimension of the SRGAP2 phenotype that our excitatory-spine imaging leaves unmeasured and that longitudinal imaging of gephyrin alongside dendritic spines would reveal.

A second body of work points to inhibition as a key component of adult structural plasticity, and is directly relevant because SRGAP2 regulates inhibitory as well as excitatory synapses. In the adult visual cortex, sensory deprivation first drives the retraction of inhibitory inputs, and the resulting reduction in inhibitory tone opens a permissive window for subsequent excitatory remodeling ^50,52^. Excitatory and inhibitory structural changes are spatially clustered along the dendrite and become more tightly coordinated by experience ^50^, and inhibitory synapses on dually innervated spines are repeatedly assembled and removed at fixed sites, providing a reversible and input-specific modulation of otherwise stable excitatory contacts ^51^. SRGAP2 sits squarely within this excitatory-inhibitory architecture, since it co-regulates the development and density of both classes of synapse and preserves the balance between them ^14,49^. Reducing SRGAP2 therefore increases inhibitory synapse density alongside excitatory synapse density, and the experience-dependent spine formation we reported in this study may be shaped by the local coordination of excitatory and inhibitory inputs.

The involvement of SRGAP2 in microglial adult function connects our findings to a broad literature on the role of microglial regulation of synapses. Microglia actively sculpt developing circuits by pruning synapses in an activity- and complement-dependent manner ^23,24^, and their physical interactions with synapses are modulated by sensory experience, with microglial contact biased toward larger dendritic spines, particularly under sensory deprivation ^25^. SRGAP2 sets the tempo of microglia maturation, and its cell-autonomous deletion produces neotenic, hyper-ramified cells that non-cell-autonomously raise spine density in adult layer 2/3 pyramidal neurons by reducing spine elimination ^16^. Our microglial-specific Srgap2-LOF cohort reproduces this stability-biased, low-elimination phenotype and extends it in two respects. First, daily imaging revealed that cell-autonomous microglial *Srgap2* deletion also permits a transient, experience-dependent increase in spine formation at the immediate onset of deprivation, a brief permissive window that returns to the wild-type pattern within a day and would be invisible at the multi-day imaging intervals used in most studies. Second, the loss of microglial SRGAP2 selectively enlarged the stable spine population, an effect present whenever SRGAP2 was reduced in microglia (*Srgap2*^+/-^ and MG-*Srgap2*^F/F^) but absent when SRGAP2 was reduced cell-autonomously in pyramidal neurons (L2/3-*Srgap2*^F/+^). This selectivity echoes the reported association between microglial contact and larger spine size under sensory deprivation ^25^, and suggests that the altered surveillance of neotenic microglia shapes the size of the spines they contact. Neuron-specific SRGAP2 haploinsifficiency drives sustained increase in spine formation following sensory deprivation in adult L2/3 PNs, whereas microglia-specific SRGAP2 deletion leads to an early but transient increased in spine formation following sensory deprivation. Both cell-type contributions seem additive in the constitutive *Srgap2*^+/-^ mice.

Several limitations of the present study should be acknowledged. Bilateral whisker trimming is a uniform sensory deprivation that lacks the competitive component of the chessboard paradigms previously used to study experience-dependent structural plasticity in adult barrel cortex ^3^, leaving open whether SRGAP2-dependent constraints on spine formation are further modulated by inter-columnar competition. Our longitudinal imaging was restricted to the apical tufts of L2/3 pyramidal neurons in layer 1 and superficial layer 2, so the basal dendrites of these neurons and the apical tuft dendritic spines of layer 5 pyramidal neurons, the canonical site of adult cortical structural plasticity, were not examined ^3,30^. Finally, our findings are confined to the S1 barrel cortex, and whether SRGAP2 reduction unmasks a comparable formation response in other cortical areas or under naturalistic experience and active learning remains to be tested ^8^.

Our results carry an evolutionary implication for the human cortex. The human-specific paralogs *SRGAP2B* and *SRGAP2C* arose through partial duplication of *SRGAP2* during the emergence of the *Homo* lineage and reduce ancestral SRGAP2A protein by ∼50% through heterodimerization and proteasome-dependent degradation ^10,12,14,15^. Therefore, partial loss of SRGAP2 function produced by human-specific SRGAP2B/C in human neurons has the same consequence as the partial loss we investigated in mice, human adult layer 2/3 pyramidal neurons may retain a heightened capacity for experience-dependent spine formation across a far longer span of adult life than other mammals. Such an effect would not merely represent a form of synaptic neoteny, originally defined as a developmental prolongation of synapse maturation ^12–14^, but maybe a persistent traits characterizing adult cortical neurons underlying heightened experience-dependent form of synaptic plasticity.

## METHODS

### Animals and Genotypes

All animal procedures were approved by the Institutional Animal Care and Use Committee (IACUC) at Columbia University. Mice were housed on a 12 h light/dark cycle with food and water available *ad libitum*. All strains were maintained in a C57BL/6J background crossed with the outbred strain 129S2/SvPasCrl (Charles River) to improve pup viability following *in utero* electroporation.

Four genotypes were used in this study. (1) C57BL/6J mice served as Wild-type controls. (2) Conditional *Srgap2* heterozygous knockout mice (*Srgap2*^F/+^) carry one copy of a floxed *Srgap2* allele ^22^. *In utero* electroporation of CAG-Cre at E15.5 drives recombination of the floxed allele specifically in targeted layer 2/3 (L2/3) pyramidal neurons, producing a cell-autonomous heterozygous loss of SRGAP2 function. (3) Constitutive *Srgap2* heterozygous knockout mice (*Srgap2*+/-) were previously described ^12^. (4) Microglia-specific *Srgap2* conditional knockout mice (MG-*Srgap2*F/F) were generated by crossing Tmem119-CreERT2 mice (The Jackson Laboratory, stock 031820) ^53^ with homozygous conditional *Srgap2* knockout mice. Microglia-specific recombination was induced by daily subcutaneous administration (3x) of 4-hydroxytamoxifen (50 μg/g body weight; TargetMol) from P3-5.

### Cohort sample sizes

The longitudinal imaging cohort comprised 19 mice across the four genotypes, contributing 264 dendritic branches, and 22,420 unique dendritic spines tracked across 8 consecutive daily two-photon imaging sessions. Per-genotype sample sizes are summarized below.

| Genotype | Mice# | Branches# | Spines# |
| --- | --- | --- | --- |
| Wild-type | 5 | 40 | 1,871 |
| *Srgap2<sup>+/-</sup>* | 4 | 51 | 7,856 |
| L2/3-*Srgap2<sup>F/+</sup>* | 5 | 110 | 5,512 |
| MG-*Srgap2<sup>F/F</sup>* | 5 | 63 | 7,181 |
| **Total** | 19 | 264 | 22,420 |

### In Utero Electroporation

*In utero* electroporation (IUE) was performed at embryonic day 15.5 (E15.5) to target L2/3 cortical pyramidal neurons, as previously described ^54^. Briefly, timed-pregnant dams were anesthetized with isoflurane, and the uterine horns were exposed through a midline abdominal incision. An endotoxin-free DNA solution was injected into the left lateral ventricle of each embryo using a heat-pulled glass micropipette and a picospritzer. The left hemisphere was consistently electroporated across all animals.

For wild-type (C57Bl6J), *Srgap2*^F/+^, and *Srgap2*^+/-^ mice, the DNA mixture contained CAG-Cre (10 ng/μl) and CAG-DIO-mGreenLantern (1 μg/μl; Addgene #164468) ^55^. The low concentration of Cre relative to the reporter ensured sparse, stochastic recombination suitable for imaging individual dendritic segments. For MG-*Srgap2*^F/F^ animals, CAG-FlpO (10 ng/μl; Addgene #60662) and FRT-mGreenLantern (1 μg/μl) were used instead, to preserve the floxed *Srgap2* alleles in labeled neurons while inducing Cre-mediated recombination in microglia-specific deletion via Tmem119-Cre^ERT2^.

Electroporation into cortical neurons was performed applying 5 pulses of 40 mV for 5 s with 100 ms intervals, using a 5 mm diameter platinum tweezer electrode and a square wave electroporator. Embryos were returned to the abdominal cavity and the incision was sutured. The dam recovered on a heated pad and gave birth naturally.

### Cranial Window Surgery

Chronic cranial windows were implanted over the left primary somatosensory cortex, barrel field (BF-S1) in mice older than P100. Mice were anesthetized with isoflurane and placed in a stereotaxic frame. The center of the barrel field was identified using stereotaxic coordinates (approximately 1.0 mm posterior and 3.0 mm lateral to bregma). A 4 mm biopsy punch was used to mark the craniotomy site on the skull, centered on BF-S1. The marked bone was removed by drilling, and a #0 round glass coverslip (4 mm diameter) was placed over the exposed cortex and secured with cyanoacrylate adhesive. A custom headplate, designed and fabricated by the Zuckerman Institute Advanced Instrumentation team, was affixed to the skull with dental adhesive cement. Mice recovered for a minimum of 3 weeks before the first imaging session to allow resolution of surgery-related inflammation.

### In Vivo Two-Photon Imaging

Longitudinal two-photon imaging was performed on a Bergamo II microscope (Thorlabs) equipped with a Coherent Chameleon Ti:Sapphire laser tuned to 920 nm for excitation of mGreenLantern. Images were acquired using ThorImageLS 4.1 (Thorlabs) through an Olympus XLPLN25XWMP2 25x water-immersion objective (1.0 NA). Z-stacks of apical dendrites from L2/3 CPNs were acquired at an XY pixel size of 0.102 μm, a z-step of 1 μm, and 50 frames averaged per plane. The number of z-planes ranged from 4 to 20 per dendrite depending on the orientation of the dendritic segments. Laser power at the sample was kept below 60 mW to minimize photodamage.

During imaging, mice were lightly anesthetized with approximately 1% isoflurane delivered via a nose cone and head-fixed using the implanted headplate. Body temperature was maintained at 37°C with a feedback-controlled thermal pad. Each imaging session lasted no longer than 1 hour per mouse.

Mice were imaged on 8 consecutive days. The same dendritic segments were relocated across sessions using vascular landmarks visible through the cranial window.

### Sensory Deprivation

Sensory deprivation was induced by bilateral whisker trimming. All whiskers on both sides of the face were trimmed to the base of the whisker pad using fine scissors under a magnified stereomicroscope, ensuring complete removal of each whisker down to the skin. Trimming began immediately after the Day 3 imaging session, such that Day 4 represented the first imaging time point under deprived conditions. Whiskers were re-trimmed daily for the remainder of the imaging period (Days 4 through 8). Days 1 - 3 served as the baseline period, and Day 3 was the final baseline session before onset of deprivation.

### Image Analysis Pipeline

Longitudinal dendritic-spine tracking was performed with a custom Python pipeline that extends our previously described RESPAN framework for automated dendritic spine segmentation ^21^, with dedicated modules for cross-session volumetric registration, 3D spine-label refinement, and longitudinal spine matching.

### Volumetric registration

For each dendritic segment, all available daily two-photon z-stacks (up to eight per segment) were co-registered to a common reference volume (the earliest imaged session, typically Day 1) using the itk-elastix library (v0.25.0). Registration proceeded in two stages: a multi-resolution rigid alignment (six-degree-of-freedom Euler transform, rotation and translation) optimizing Mattes mutual information, followed by a non-rigid B-spline refinement stage (final grid spacing 8.0 × 8.0 × 4.0 μm in X, Y, and Z) optimizing normalized cross-correlation, with a transform-bending-energy penalty (weight 1.0) to suppress residual local distortions. The anisotropic voxel geometry of the acquisition (XY = 0.102 μm; Z = 1.0 μm) was respected throughout by an anisotropic multi-resolution pyramid schedule whose per-dimension downsampling factors were chosen such that each spatial dimension retained at least four voxels at the coarsest level; the number of resolution levels was configured at four but automatically capped per volume to satisfy this constraint (effective range across the dataset, 1 to 3 levels). Each stage drew 2048 spatial samples per iteration (resampled at every iteration) with a minimum valid-sample fraction of 0.05 and ran for up to 500 iterations. Registration was initialized by geometric-center alignment; if elastix raised a convergence error, the pipeline automatically retried rigid-only registration with geometric-center initialization, and, as a final fallback, rigid-only registration with origin-based (identity) initialization. The fitted transform was then propagated to the auxiliary per-session channels (spine label maps, dendrite masks, dendrite skeletons, and along-dendrite arc-length fields) with nearest-neighbor interpolation for categorical and coordinate channels and linear interpolation for fluorescence. Registration quality was quantified for each moving session by computing the normalized mutual information (NMI) and the normalized cross-correlation (NCC) between the final registered moving volume and the reference volume.

### Spine segmentation

Prior to registration, spines and dendrites were segmented on each raw two-photon volume using RESPAN ^21^. Per-session outputs comprised a 3D connected-component label map of individual dendritic spines (one integer label per spine instance), a binary dendrite mask, a single-voxel-wide dendrite skeleton, and a scalar along-dendrite arc-length field encoding geodesic distance along the skeleton from a fixed endpoint.

### 3D spine-label refinement

To resolve occasional RESPAN false merges in which two closely apposed spines received a single connected-component label, each spine label was further refined in 3D. The original spine label mask was eroded using a planar 3×3 structuring element applied in XY only (implemented as a (3,3,3) kernel with only the central Z-plane active), intentionally skipping the low-resolution Z axis to preserve thin protrusions. Connected components of the eroded mask were identified with 6-connectivity and filtered by retaining only components of at least 10 voxels. Remaining components whose physical centroids fell within 0.5 μm of one another were then iteratively merged by collapsing the higher-indexed seed into its lower-indexed neighbor, until no pairs remained within threshold. When multiple valid seeds survived within an original label, an anisotropy-aware 3D Euclidean distance transform was computed over the original mask, and the mask was split into per-seed regions via marker-controlled watershed. False-split fusion (the merging of two adjacent RESPAN labels into a single spine) was not performed.

### Longitudinal spine matching

Individual spines were tracked across sessions by constructing, per dendritic segment, a spine-tracking graph in which every spine received a persistent global identifier. Correspondences between the reference session (typically Day 1) and every subsequent session were computed by formulating spine-to-spine matching as a bipartite assignment problem and solving it with the Hungarian algorithm, using the 3D intersection-over-union (IoU) between pairs of registered spine label volumes as the cost. After assignment, candidate matches were independently gated by three rejection criteria: (i) IoU ≥ 0.05; (ii) 3D physical centroid-to-centroid distance ≤ 1.5 μm; and (iii) volume ratio. Spines on session N that failed to be assigned to a reference-session anchor were re-linked in two further passes: a consecutive-day pass (Day N matched to anchors on Day N−1) and, for spines still unmatched, a one-day-gap pass (Day N matched to anchors on Day N−2), rescuing spine identities lost to cumulative registration drift or transient imaging artifacts without creating new global identifiers. All downstream turnover and density analyses were performed directly on the resulting tracking table; no gap imputation was applied.

### Common dendritic ROI and classification of spine dynamics

For each dendritic segment, a common dendritic ROI was defined to ensure that longitudinal turnover statistics were computed over an identical stretch of dendrite at every time point. Each spine centroid was mapped into a one-dimensional position along the reference-day dendrite by evaluating the RESPAN along-dendrite arc-length field (geodesic distance along the skeleton from a fixed endpoint) at the nearest reference-day dendrite voxel. For each session, the range of this coordinate attained by the registered dendrite mask was computed (subsampling up to 200 dendrite voxels per session to control runtime). The common ROI was then defined as the intersection of these ranges across all available imaging sessions, taken as the maximum of the per-session lower bounds and the minimum of the per-session upper bounds. For dendritic segments in which branching was detected, this intersection was computed per branch, and each spine was assigned to its enclosing branch. Spines whose reference-frame dendrite coordinate fell outside the common ROI were flagged and excluded from per-day-pair composition analyses. Within the common ROI, and for each consecutive day pair (Day N−1 → Day N), each spine was classified as stable if its global identifier was present on both Day N−1 and Day N, as formed on Day N if its global identifier was present on Day N but absent on Day N−1, and as eliminated on Day N if its global identifier was present on Day N−1 but absent on Day N. Spines absent from both days of the pair were excluded from the per-day-pair composition.

### Spine density and per-day-pair spine composition

For each dendritic segment, longitudinal summary statistics were computed from the tracking table. Spine density on session N was computed per dendritic segment from that session’s 3D spine and dendrite segmentation, as the total number of spine labels on session N divided by the total dendritic length of that segmented ROI on session N (reported in spines/μm).

Per-day-pair spine composition was computed for each consecutive day pair (Day N−1 → Day N) and restricted to the common dendritic ROI defined above. For each dendritic branch, every spine present on Day N−1, Day N, or both was classified into one of three categories based on its cross-session presence: stable spines, present on both days; newly formed spines, present on Day N but absent on Day N−1; and eliminated spines, present on Day N−1 but absent on Day N. Spines absent from both days of the pair were excluded from the per-day-pair analysis. Let *n*_stable_, *n*_formed_, and *n*_eliminated_ denote the corresponding spine counts within the common ROI of a branch at a given day pair. Per-branch fractions of stable, formed, and eliminated spines were defined as *n*_stable_/*D*, *n*formed/*D*, and *n*_eliminated_/*D*, respectively, where *D* = *n*_stable_ + *n*_formed_ + *n*_eliminated_. By construction, the three fractions sum to one per branch per day pair, and each metric is reported on the second day of the pair (Day N). Aggregation across branches, segments, animals, and genotypes is described in statistical analysis section.

### Pipeline validation

Cross-session registration quality was quantified for each moving-session/reference-session pair by the normalized cross-correlation (NCC) between the registered moving volume and the Day 1 reference volume. Registration increased the median per-pair NCC from approximately 0.31 to approximately 0.80 across the full dataset, with a significant Before-vs-After improvement within every genotype (two-way ANOVA, factors Genotype × Registration state; p < 0.0001 within each genotype; **Suppl. Fig. 1c-d**). Moving-session/reference-session pairs with post-registration NCC below 0.6 were excluded from downstream longitudinal analyses.

Longitudinal tracking precision was quantified by the residual centroid displacement between consecutive registered sessions for spines whose persistent global identifier was preserved across the pair. The median per-spine centroid displacement was 0.499 μm and 92.6% of matched-spine displacements fell below 1.0 μm (**Suppl. Fig. 1e**), consistent with previously reported inter-session spine localization precision for chronic *in vivo* two-photon imaging ^2,3,5^.

### Statistical Analysis

Statistical analyses were performed in Python or PRISM. Sample sizes refer to the unit of analysis appropriate to each measurement: individual dendritic segments for spine density and inter-spine distance comparisons; individual spine-centroid displacements for tracking-quality summaries; and per-genotype paired volume pairs for registration-quality comparisons. For Day 1 cross-sectional analyses (**Fig. 3c**, **3d**), all dendritic segments imaged at the Day 1 baseline were pooled per genotype regardless of subsequent longitudinal retention (Day 1 segment n: WT, 79; *Srgap2*^+/-^, 71; L2/3-*Srgap2*^F/+^, 135; MG-*Srgap2*^F/F^, 218); all longitudinal analyses (**Fig. 3e**, **Fig. 4**) are restricted to the longitudinal-tracked subset summarized in Animals and Genotypes section above. Per-genotype sample sizes (mice, segments, branches, spines, and spine-pair observations) are reported in Animals and Genotypes section above and summarized in the corresponding figure legends.

Tests specific to each figure panel are as follows. **Suppl. Fig. 1c,d** (registration NCC): normalized cross-correlation values before and after registration were compared per genotype across all consecutive-day volume pairs; post-registration distributions across genotypes (Suppl. Fig. 1c) were summarized descriptively. **Fig. 3c** (Day 1 spine density): spine density across the four genotypes was compared with a Kruskal-Wallis test; each *Srgap2*-LOF genotype was then compared pairwise to Wild-type using Dunn’s multiple comparisons test, with adjusted p values reported. **Fig. 3d** (inter-spine distance distributions): the cumulative distribution of inter-spine distances for each *Srgap2*-LOF genotype was compared to Wild-type using a two-sample Kolmogorov-Smirnov test; reported p values are unadjusted across the three Wild-type-referenced contrasts. **Fig. 4b-d** (per-day-pair spine composition): at each consecutive day pair (Day pairs 1-2 through 7-8), the per-branch fraction of spines classified as stable (b), newly formed (c), or eliminated (d) was compared between each Srgap2-LOF genotype and Wild-type at the same day pair using a two-sided Mann-Whitney U test.

Statistical significance thresholds are indicated throughout the figures as n.s. (p ≥ 0.05), *p < 0.05, p < 0.01,* p < 0.001, and ****p < 0.0001; exact p values and test statistics are reported in the corresponding figure legends where feasible.

## Supplementary figures

**Supplementary Figure 1.**
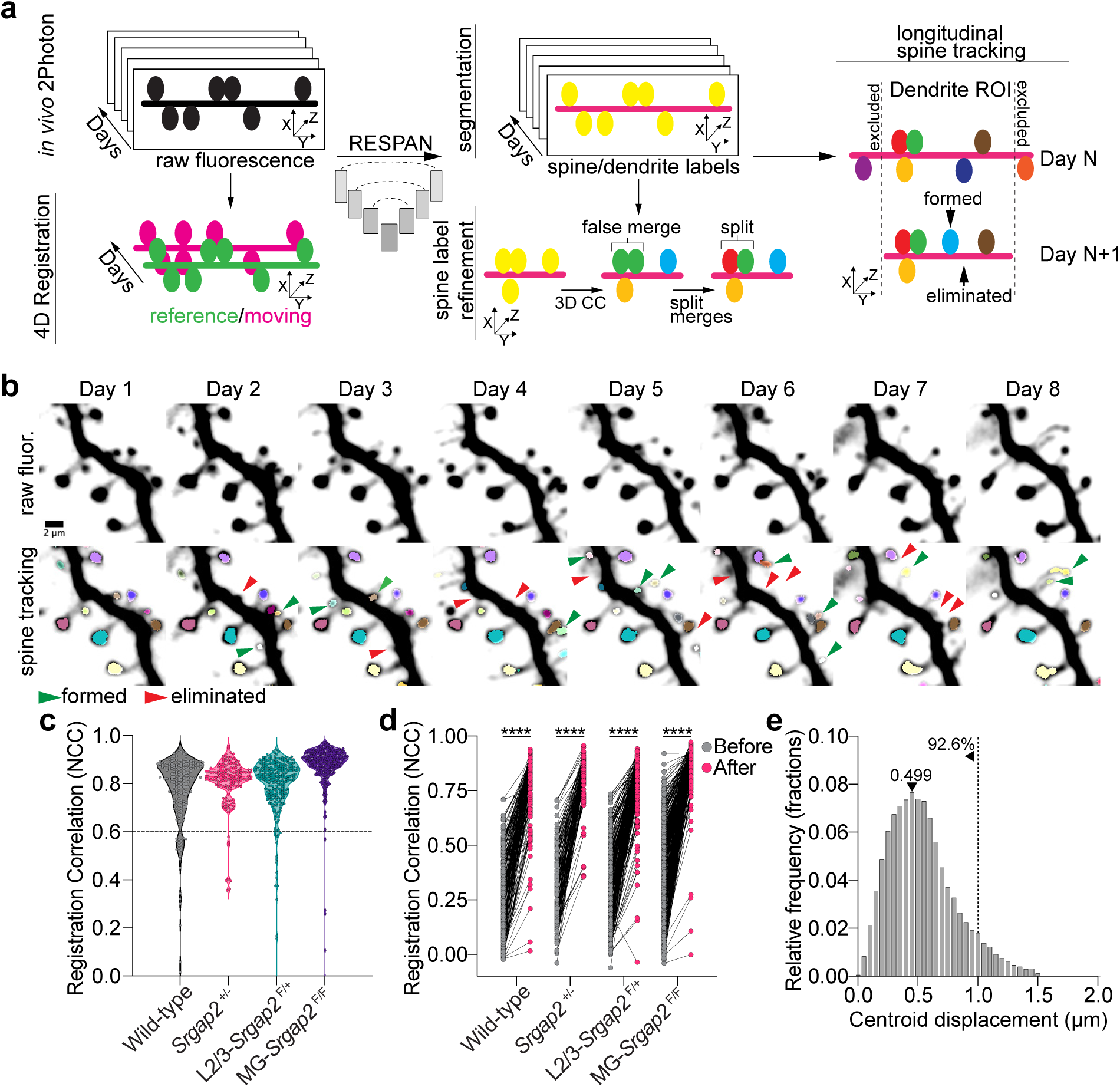
Deep-learning based analysis pipeline enables reliable and unbiased longitudinal spine tracking. (**a**) Analysis pipeline overview. Daily *in vivo* two-photon z-stacks acquired across eight consecutive imaging sessions are independently (i) segmented on the raw volumes using RESPAN (producing per-session spine label maps, dendrite masks, single-voxel-wide skeletons, and along-dendrite arc-length fields) and (ii) co-registered in 3D to a common reference volume (typically Day 1) via a multi-resolution rigid plus non-rigid B-spline transform (itk-elastix); the reference volume is shown in green and the moving volume in magenta. The fitted transform is then propagated to all RESPAN outputs with nearest-neighbor interpolation, bringing every per-session channel into the reference-day coordinate frame. Spine labels are further refined in 3D to resolve occasional RESPAN false merges by seed-based splitting of closely apposed spines sharing a single connected-component label (3D CC). Longitudinal spine tracking is performed in 3D by matching registered spine label volumes from each subsequent session to the reference session via the Hungarian algorithm on intersection-over-union, with rescue passes for unmatched spines against the immediately preceding and the once-removed preceding session. A common dendritic ROI, defined as the dendrite extent shared across all imaging sessions (dashed vertical lines), ensures that the same stretch of dendrite is analyzed at every time point; spines outside the common ROI are flagged and excluded. Each tracked spine is then classified as persistent, formed, or eliminated relative to the previous session. (**b)** Example of automated spine tracking across eight daily sessions for a single dendritic segment. Top row: raw fluorescence maximum-intensity projections. Bottom row: RESPAN segmentation overlaid on raw fluorescence with longitudinal tracking; each tracked spine retains a consistent color reflecting its persistent global identifier across sessions. Arrows indicate formed spines (green) and eliminated spines (red). Scale bar, 2 μm. (**c**) Registration quality across genotypes. Violin plots of normalized cross-correlation (NCC) between each registered moving session and the reference session (typically Day 1), per dendritic segment, for Wild-type, *Srgap2*^+/-^, L2/3-*Srgap2*^F/+^, and MG-*Srgap2*^F/F^. Each point is one moving-session/reference-session pair. The dashed line at NCC = 0.6 indicates the registration-quality threshold; segments whose post-registration NCC fell below 0.6 were excluded from downstream longitudinal spine tracking. (**d**) Registration quality before and after volumetric registration, per genotype. Paired plot of NCC values computed on the same moving-session/reference-session pairs as in (c), before (gray) and after (magenta) applying the fitted transform. Two-way ANOVA with factors Genotype and Registration state (Before, After); ****P < 0.0001 for the Before-vs-After contrast within every genotype. **e** | Distribution of per-spine centroid displacements between consecutive registered sessions (Day N to Day N+1), pooled across all genotypes, for spines whose global identifier was preserved across the pair. Median displacement = 0.499 μm; 92.6% of matched-spine displacements fall below 1.0 μm (dashed line).

**Supplementary Figure 2.**
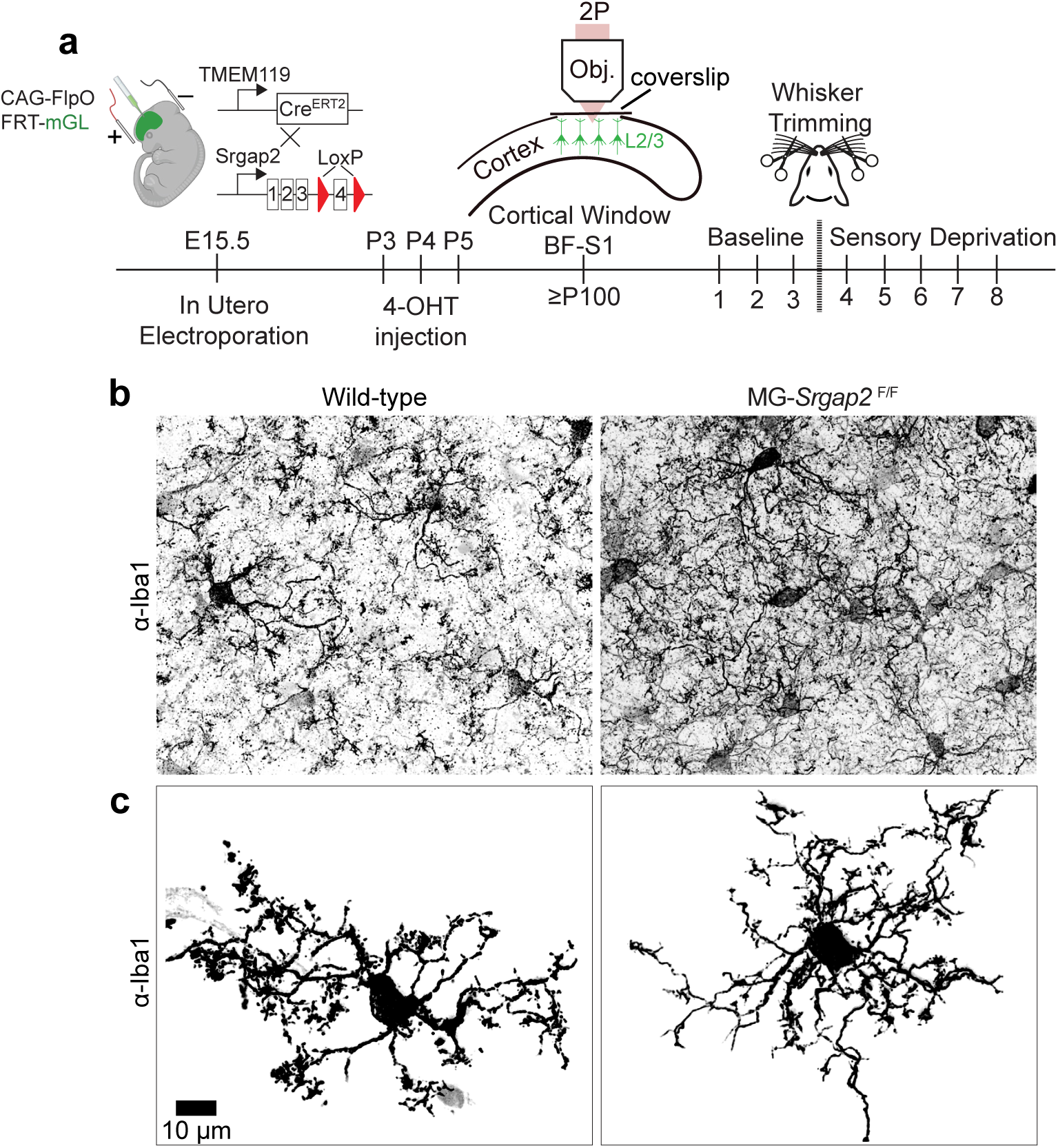
Microglia-specific *Srgap2* deletion re-produces a hyperramified microglial morphology, confirming functional Cre recombination. (**a**) Cell-type-specific genetic strategy. Layer 2/3 pyramidal neurons were sparsely labeled by *in utero* electroporation at E15.5 (CAG-FlpO/FRT-mGreenLantern) in TMEM119-CreERT2;*Srgap2*^F/F^ mice. 4-hydroxytamoxifen (4-OHT) was administered daily from P3-P5 to induce Cre-mediated *Srgap2* excision selectively in all microglia. A chronic cranial window was implanted over barrel field somatosensory cortex (BF-S1) at ≥P100 and dendritic spines were imaged across 8 daily sessions (Baseline, Days 1-3; Sensory Deprivation, Days 4-8). (**b**) Tiled α-Iba1 immunostaining of BF-S1 from a WT animal (left) and an MG-*Srgap2*^F/F^ animal (right). Both animals were drawn from the main longitudinal imaging cohort; tissue was collected after terminal Day 8 imaging. (**c**) Higher-magnification α-Iba1 images of individual microglia from the same sections as in **b**. Microglia in MG-*Srgap2*^F/F^ are hyperramified relative to WT, consistent with functional loss of SRGAP2 in microglia after Cre activation. Scale bar, 10 μm.

**Supplementary Figure 3.**
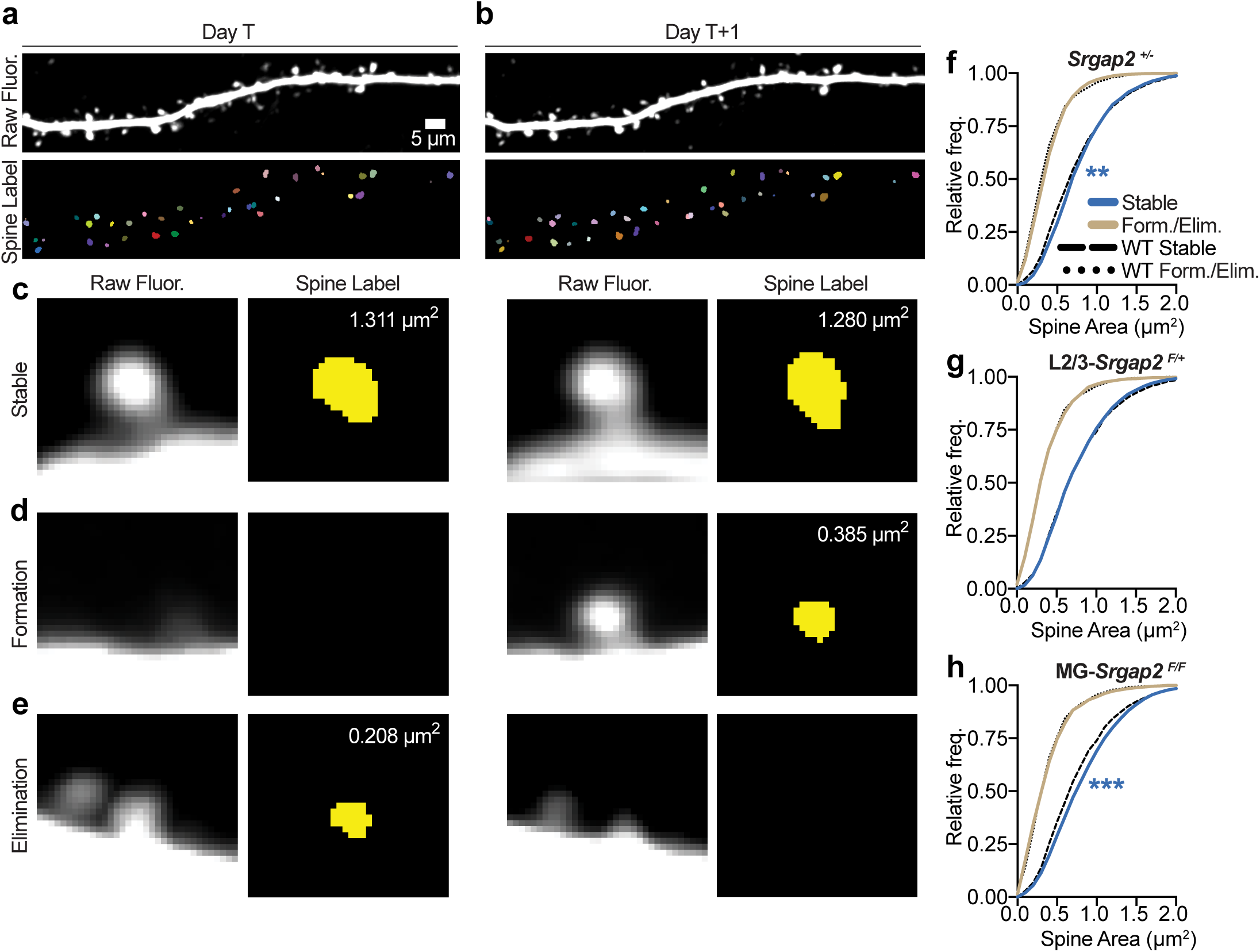
Stable spines are larger than transient spines across all genotypes, and microglial *Srgap2* reduction selectively enlarges the stable spine population. (**a-b**) Example dendritic segment imaged on two consecutive daily sessions, Day T (a) and Day T+1 (b), each shown as the raw two-photon maximum-intensity projection (Raw Fluor., top) and the corresponding RESPAN spine-label map (Spine Label, bottom), in which individual segmented spines are shown as distinctly colored labels. Spine area was measured as the 2D projected area of each spine-label mask on the maximum-intensity projection. Scale bar, 5 μm (a, applies to b). (**c-e**) Representative single spines illustrating the three longitudinal fate classes, each shown as raw fluorescence and the corresponding spine-label mask at Day T and Day T+1, with the projected spine area (μm²) indicated. **c** | A stable spine, present on both days (1.311 μm² at Day T, 1.280 μm² at Day T+1). (**d**) A newly formed spine, absent at Day T and present at Day T+1 (0.385 μm²). (**e**) An eliminated spine, present at Day T (0.208 μm²) and absent at Day T+1. (**f-h**) Cumulative distributions of single-spine area for spines observed during the pre-deprivation baseline epoch (Days 1-3), comparing stable spines (blue) and transient spines (newly formed and eliminated, pooled; tan, labeled Form./Elim.) for each *Srgap2*-loss-of-function genotype against the corresponding wild-type reference distributions (WT Stable, dashed black; WT Form./Elim., dotted black). In every genotype, stable spines are approximately two-fold larger than transient spines. Panel f: *Srgap2*^+/-^. The stable spine population is significantly right-shifted relative to WT stable spines (median area 0.749 vs 0.697 μm²; D = 0.069, **P = 0.003); the transient population does not differ from WT. Panel g: L2/3-*Srgap2*^F/+^. Neither the stable nor the transient population differs from WT (stable median 0.697 vs 0.697 μm²; n.s.). Panel h: MG-*Srgap2*^F/F^. The stable spine population is significantly right-shifted relative to WT stable spines (median area 0.801 vs 0.697 μm²; D = 0.080, ***P = 0.0003); the transient population does not differ from WT. Two-sample Kolmogorov-Smirnov test of each genotype population against the corresponding WT population. Baseline-epoch (Days 1-3) spines from the same longitudinal cohort as Figure 4 (19 mice, 264 branches). n (stable, transient spines): *Srgap2*^+/-^ 3,034, 2,297 (f); L2/3-*Srgap2*^F/+^ 2,195, 1,407 (g); MG-*Srgap2*^F/F^ 3,530, 1,651 (h); wild-type reference 858, 568 (f-h). n.s., not significant; **P < 0.01, ***P < 0.001.

